# Interior of sand fly (Diptera: Psychodidae) abdomen reveals novel structures involved in pheromone release: discovering the Manifold

**DOI:** 10.1101/2021.08.18.456824

**Authors:** G.B. Tonelli, J.D. Andrade-Filho, A.M. Campos, C. Margonari, A.R. Amaral, P. Volf, E. Shaw, J.G.C. Hamilton

## Abstract

The males of many species of New World Phlebotomines produce volatile terpenoid chemicals which have been shown in *Lutzomyia longipalpis* s.l. and *L. cruciata* to be sex/aggregation pheromones which attract female and male conspecifics. Pheromone is produced in secretory cells surrounding a cuticular reservoir which collects the pheromone and passes it through a cuticular duct to the surface of the insect. On the surface the pheromone passes through a specialised structure prior to evaporation. The shape and distribution of the structures are highly diverse and differ according to species. They range in appearance from slightly raised domes (papules) to almost spherical apple shaped structures to slight depressions with central spikes and all with a central pore. They can occur either singly or in many hundreds distributed on most abdominal tergites or grouped on one. The pheromone secreting apparatus in sand flies and other insects have historically been examined from the exterior using scanning electron microscopy (SEM) and from the interior using transmission electron microscopy. In this study we used SEM to examine the interior cuticular structure of 3 members of the *Lutzomyia longipalpis* s.l. species complex and *Migonemyia migonei* and found a new structure associated with pheromone release which we have called the Manifold. The Manifold is a substantial structure siting in-line between the cuticular duct and the underside of the tergite. Differences in the size and shape of the Manifold may be related to the chemical structure of the pheromone. In addition to the importance of this hitherto unknown structure in the production, dissemination and ecology of the pheromone, as well as its potential taxonomic value, examination of the interior cuticle by SEM may help locate the secretory apparatus in important vector species where pheromonal activity has been inferred from behavioural studies but the external secretory structures or potential pheromones have not been found.

## INTRODUCTION

Members of the sand fly species complex, *Lutzomyia longipalpis* s.l. (Lutz and Neiva, 1912) have been identified as vectors of the Protist parasite *Leishmania infantum* Nicolle, 1908, the etiological agent of visceral leishmaniasis (VL) (Deane 1956; Deane and Deane 1954; Lainson and Rangel 2005). There is a close relationship between the distribution of VL cases and the distribution of *L. longipalpis* s.l. throughout most of Brazil and it has been proposed that the urbanization of members of this complex and it’s anthropophilic behaviour have increased the incidence of VL in many Brazilian states (Casanova *et al.* 2015; Rangel and Vilela 2008).

In some regions of Brazil, where *L. longipalpis* is not the most abundant sand fly species, VL cases are associated with another incriminated vector, *Migonemyia migonei* (França, 1920) (de Carvalho *et al.* 2010; Rangel and Lainson 2009). This species is considered to be a potential vector because of its’ distribution, prevalence, anthropophily and the detection of *Leishmania* DNA in blood-fed females (de Carvalho *et al.* 2010). In addition, based on evidence of the development of late-stage parasite forms in artificially infected sand flies this species is considered permissive for transmission of *Le. infantum* (Guimarães *et al.* 2016). *Migonemyia migonei* has also been implicated as a vector of *Le*. (*V*.) *braziliensis*, the etiological agent of cutaneous leishmaniasis in different Brazilian regions (de Pita-Pereira *et al.* 2005; de Queiroz *et al.* 1994).

Male *L. longipalpis* s.l. produce sex/aggregation pheromones which when present with host odour are attractive to conspecific males (Morton and Ward 1989; Nigam and Ward 1991; Ward *et al.* 1991) and lead to the formation of leks on or near host animals where the males compete with each other for access to mating opportunities (Jones and Hamilton 1998; Morrison *et al.* 1995). Females are attracted by the combination of the male produced pheromone and host odour (Bray and Hamilton 2007; Kelly and Dye 1997; Spiegel *et al.* 2016). They arrive at the lekking site after the males (Kelly and Dye 1997), choose a mate (Jones and Hamilton 1998) take a blood-meal and depart (Ward *et al.* 1993). It has been proposed that synthetic sex/aggregation pheromone co-located with insecticide could be used for vector control (Bray *et al.* 2009; Bray *et al.* 2014; Courtenay *et al.* 2019). In addition, analysis of these pheromones has also been used as a taxonomic tool (Foster and Dugdale 1988) to differentiate between individual members of the *L. longipalpis* species complex (Hamilton *et al.* 2005; Hickner *et al.* 2021; Souza *et al.* 2017; Spiegel *et al.* 2016; Vigoder *et al.* 2020).

In *L. longipalpis* s.l. the sex/aggregation pheromone has been associated with cuticular structures on the surface of tergites III or III and IV and associated with underlying glandular tissue where they have the visual appearance of either 1 or 2 pale spots (1S or 2S)(Boufana 1990; Spiegel *et al.* 2016). Under SEM the external cuticular structures appear as small round elevations (papules) with a central pore (mean diameter 0.25 μm) and a density of 21±2 (1S) or 19±2 (2S) μm^−2^ (Lane and Ward 1984; Spiegel *et al.* 2005). The pheromone is believed to be produced by the glandular cells that underly the papules and to be passively transported to the surface pores via a cuticular duct (Boufana 1990; Spiegel *et al.* 2002). The sex/aggregation pheromones are different in each member of the complex (Hamilton *et al.* 2005) and 5 members can be distinguished two have been characterised as methylsesquiterpene (C16), two as diterpene (C20) (Hamilton *et al.* 2004; Palframan *et al.* 2018; Spiegel *et al.* 2016). Those *L. longipalpis* that produce the methylsesquiterpene, (*S*)-9-methylgermcrene-B, can be further subdivided into 2 population types represented by those from Sobral (CE) and Lapinha (MG) (Hamilton *et al.* 2005; Hickner *et al.* 2021).

Possible pheromone associated tergal structures have also been observed in other sand fly species where they occur in a variety of forms (Ward *et al.* 1991; Ward *et al.* 1993). For example, in *Evandromyia lenti* and *E. carmelinoi*, apple-shaped structures (2.7±0.1 μm in diameter x 1.6±0.1 μm in height and 2.5±0.2 μm in diameter and 1.2±0.2 μm in height respectively) with a central pore are present on the V and VI tergal segments (Spiegel *et al.* 2002). Terpenoids including oxygenated compounds are produced in some of these other species, but they do not appear to be present in all those that have structures (Hamilton *et al.* 2002; Hamilton *et al.* 1999; Serrano *et al.* 2016) and behavioural evidence for pheromonal activity for any of these compounds is lacking. SEM analysis of the tergal structures in *M. migonei* have revealed that they form of a shallow crater (average diameter ca. 3.2 μm) with a central pit (av. diameter ca. 0.4 μm) containing a central spike (height ca. 0.2 μm) within it (Ward *et al.* 1991; Ward *et al.* 1993). There is some behavioural evidence that this species also produces a sex pheromone (Costa 2016).

Although TEM studies have also indicated the presence of internal cuticular structure e.g. the end apparatus and cuticular duct associated with the secretory cells (Boufana 1990; Noirot and Quennedey 1974; Spiegel *et al.* 2002), there has been no SEM investigation of these internal cuticular structures in sand flies or as far as we can determine any other insect group. Therefore, this SEM study was undertaken to investigate the internal cuticular structures associated with pheromone production and release and to compare the morphology of these structures in three members of the *L. longipalpis* species complex and *M. migonei*.

## MATERIAL AND METHODS

### Sand flies

The male *L. longipalpis* used in the study were obtained from colonies held at Lancaster University, UK and the *M. migonei* were obtained from a colony held at Charles University, Czech Republic. The *L. longipalpis* colonies were representative of 3 of the 5 pheromone types found in Brazil (Hamilton *et al.* 2004; Hamilton *et al.* 2005; Hickner *et al.* 2021) and were established from females originally collected using miniature CDC light traps in chicken shelters (Table 1). The *M. migonei* colony was also established from material originally collected using CDC light traps in Baturité, Ceará, Brazil (04°19’41”S, 38°53’05”W). The *L. longipalpis* colonies were maintained in an insectary (28±2°C, 80±5% RH and a 12:12 light:dark (L:D) photoperiod) and all males used in this study were 7d old and classified as two spot (2S) (Mangabeira Filho 1969). The *M. migonei* colony was maintained under slightly different conditions (Volf and Volfova 2011) and males used were 5-7d old.)

**Table 1.** Original collection site and pheromone type of the members of the *L. longipalpis species* complex held at Lancaster University used in the study.

| collection locality | grid reference | pheromone type |
| --- | --- | --- |
| Campo Grande - MS | 20° 28'S, 54° 37'W | (S)-9-methylgermacrene-B (9MGB) |
| Jacobina - BA | 11° 11'S, 40° 31'W | 3-methyl-α-himachalene (3MH) |
| Sobral - CE | 3° 41'S, 40° 20'W | Sobralene (SOB) |

The male sand flies used in this study were removed from the colony and killed by placing them in a freezer (−5°C) for 20 mins. They were then placed in a plastic screwcap vial and covered with a few drops of ethanol (70%) and stored (−20°C) until dissection.

### Dissection

To prepare the male sand fly abdomen for SEM, a male was placed in a drop of saline solution (1% w/v) on a glass microscope slide. The entire abdomen was removed from the thorax and the entire abdomen or tergites III and IV were excised from the other abdominal segments with entomological needles under a dissecting microscope (Stemi 508, Carl Zeiss Ltd, Cambridge, UK). The interior of the whole abdomen or the abdominal segments III and IV were then exposed by a further dorsoventral incision.

### Digestion, cleaning and drying of cuticle sections

To remove the internal soft tissue covering the interior cuticular structures we submerged the dissected abdominal samples in KOH in glass Petri dishes placed on a plate rocker. *L. longipalpis* samples were digested in KOH (10% w/v) for 4 hours and *M. migonei* were digested in KOH (10% w/v) for 24 hours. After the KOH digestion, the samples were washed in saline solution (1% w/v) in a Petri dish for 5 min (3 times) followed by a final rinse in distilled water. The samples were then dehydrated by washing in alcohol (50%, 70%, 90% and 100%) for 5 min each and then left overnight in a fume hood in hexamethyldisilazane until completely dry.

### Scanning Electron Microscopy (SEM)

After the digestion, cleaning and drying, samples were mounted on SEM stubs with double sided adhesive tape and sputter coated with gold (20nm) (Edwards S150A; Edwards UK, Burgess Hill, UK). The samples were then examined with a scanning electron microscope (JEOL JSM-7800F and JEOL JSM-5600; Jeol (UK) Ltd, Welwyn Garden City, UK) operated at 18kV. In total four Campo Grande, three Sobral and three Jacobina *L. longipalpis* specimens as well as four *M. migonei* specimens were prepared and examined by SEM. Digital images of 10 randomly selected manifold structures from each sand fly specimen were made and the dimensions of the observed structures were measured using Image J®software.

### Measurements of the pheromone gland structures

We measured sizes of 4 elements of the secretory apparatus; width, height, reservoir + cuticular duct length and secretory apparatus length. We measured the sizes of 10 of each of the 4 structures from five different specimens of each of the three *L. longipalpis* pheromone types (total measurements = 600).

Comparison of the size of the different pheromone gland structures measured in each of the three *L. longipalpis* chemotypes and *M. migonei* was made by Generalized Linear Model (GLM). We assumed that there was no difference in the morphology of the structures of individuals from the same location. The measurements of each part of the structure measured were used as response variables, while the colonies were considered to be explanatory variables. Tukey’s test was used to determine which measurement were different from each other.

All the models were made using R (v3.6.1, R Development Core Team 2016), following by residual analysis to standardize the data distribution.

### Ethics Statement

Sand fly blood feeding at Lancaster University for colony maintenance was performed according to the guidelines and regulations of the Animals in Science Regulation Unit (ASRU) and in accordance with the terms of a regulated licence (PPL P2DB5013A) and in compliance with the Animals (Scientific Procedures) Act (ASPA) 1986 (amended 2012) regulations and was consistent with UK Animal Welfare Act 2006. All procedures involving animals were reviewed and approved by the Faculty of Health and Medicine Ethical Review Committee (FHMREC15125) at Lancaster University. Sand fly blood feeding at Charles University for colony maintenance was performed in accordance with institutional guidelines and Czech legislation (Act No. 246/1992, amendment No. 359/2012) which complies with relevant European Union guidelines for experimental animals. All procedures involving animals were approved by the Committee on the Ethics of Laboratory Experiments of the Charles University (Registration Number: MSMT-8604/2019-6).

## RESULTS

### The *Lutzomyia longipalpis* complex secretory apparatus

Preliminary investigation showed that digestion of sand fly samples in KOH (5% and 10%) over 4 hours removed tissue from the inner cuticular surface of the abdomen without damaging the target structure.

Examination of the interior surface of the *L. longipalpis* abdomen showed structures that were distributed over an area that matched both the size and shape of the pale spots previously observed on the external surface of tergites III and IV (Lane and S. 1990; Spiegel *et al.* 2002; Ward *et al.* 1993) Fig. 1.

**Fig. 1.**
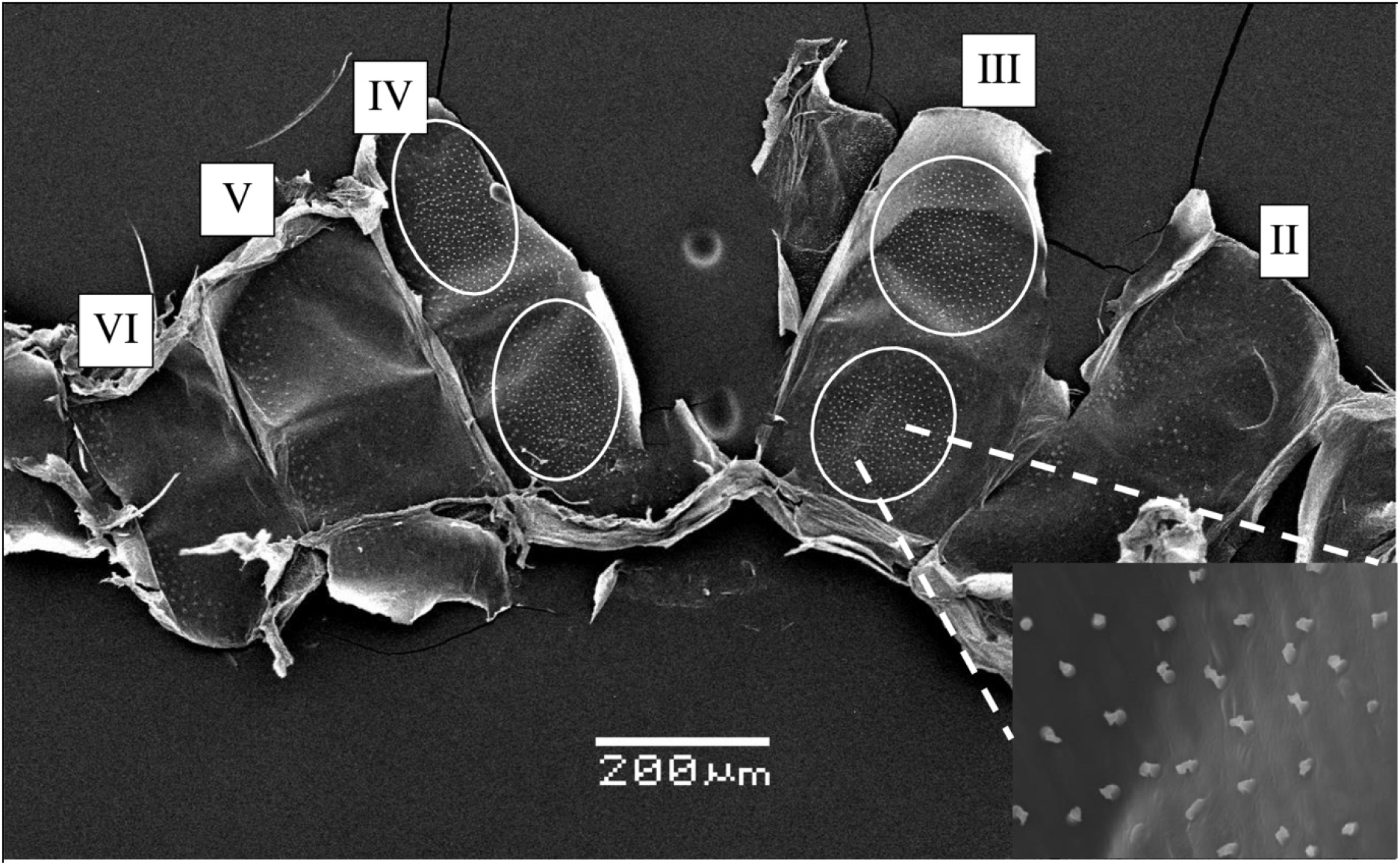
SEM of the interior cuticular surface of abdominal segments II-VI of *L. longipalpis* from Campo Grande showing the areas corresponding to pale patches normally seen from the exterior. Tergites II to VI are indicated by Roman numerals. The areas of the internal surface corresponding to the pale spots seen from the exterior area indicated by the white oval shapes. The insert is a close-up magnification of the end apparatus and associated cuticular structures seen within the oval-shaped (pale patch) areas.

Density of these structures in the samples from Campo Grande was approximately 13/1000 μm^2^ (ca. 1627 structures in total), Jacobina 18/1000 μm^2^ (ca. 1415 structures in total) and Sobral 18/1000 μm^2^ (ca. 3469 structures in total).

Observation of the morphology of the internal cuticular structure which remained after KOH digestion indicated that two sections were present; the first was a section which connected to the interior wall of the tergite (or which is an extension of the tergite) and which we have called the manifold (Fig. 2A). The manifold has two distinct parts; the base and a distally positioned section, the ring, which has the appearance of a doughnut shaped thicker ring of cuticle (Fig. 2B). The second part of the whole structure is the cuticular duct (chitinous duct (Lane and S. 1990)) which is connected to the manifold at the proximal end and which terminates in the secretory reservoir at the distal end (Fig. 2C). The secretory reservoir is seen to be a cuticular bag that can assume different shapes. Both the cuticular duct and the secretory reservoir are structures that have been previously observed in TEM studies (Boufana 1990; Spiegel *et al.* 2002) but have not been observed in SEM studies. All parts can together be described as the secretory apparatus (Fig. 2D). In some cases, during the preparation of the samples the ductule/reservoir complex become detached from the manifold structure showing that the interior of the manifold appears to be hollow.

**Fig. 2.**
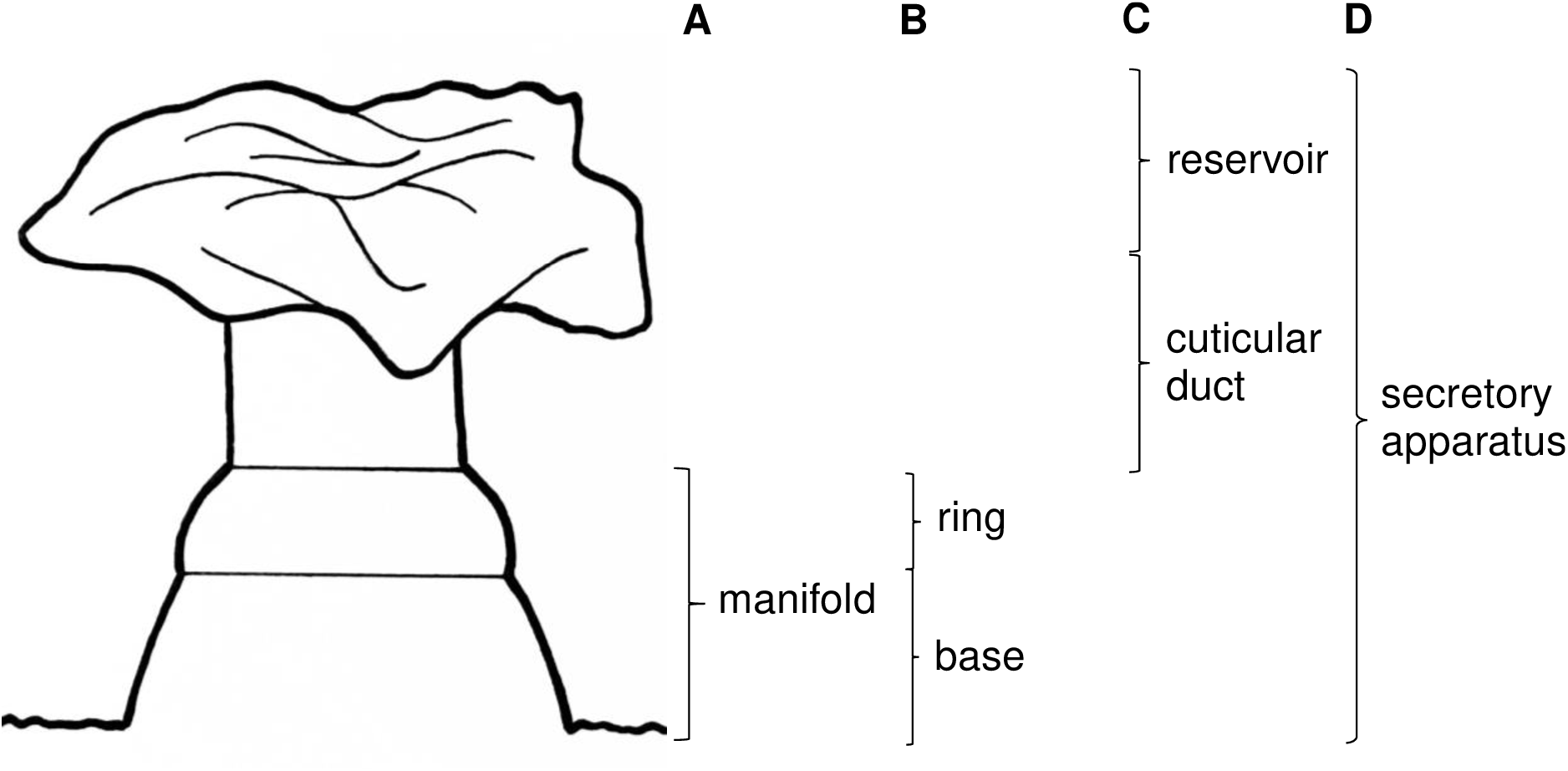
Drawing of the components of the secretory apparatus of *Lutzomyia longipalpis* from Campo Grande, Brazil. **A)** manifold connected to the inner surface of the abdominal cuticle; **B)** components of the manifold, ring + base; **C)** secretory reservoir + cuticular duct; **D)** secretory apparatus, reservoir + cuticular duct + manifold.

The secretory apparatus of the three members of the *L. longipalpis* complex examined in this study are shown in Fig. 3.

**Fig. 3.**
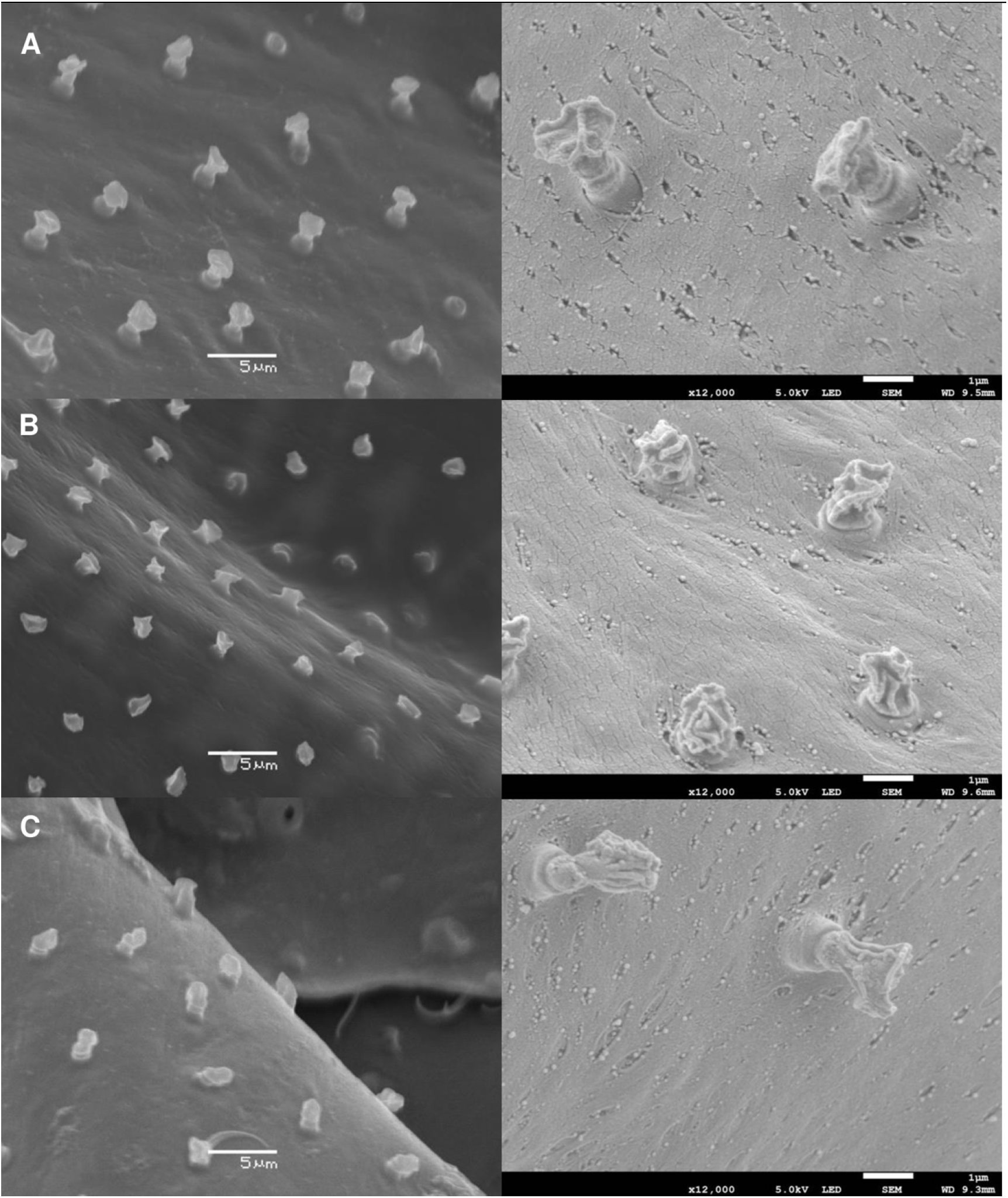
SEM images of the inner cuticle surface of the abdominal tergites of 3 members of the *L. longipalpis* s.l. species complex showing the cuticular elements; manifold, reservoir and cuticular duct, of the secretory apparatus. Secretory apparatus observed by SEM after KOH digestion of *L. longipalpis* abdominal tergites from; **A)** Campo Grande, **B)** Sobral and **C)** Jacobina. Images on the left side (x3,500 magnification) were taken on a Jeol JSM-5600. Images on the right (x12,000 magnification) were taken on a Jeol JSM-7800F.

There was a highly significant difference in the widths of the Manifolds (Fig. 2) of the 3 types of *L. longipalpis* (df=147; F=15.17; *P*<0.001). The Campo Grande manifold was significantly wider (mean±sem; 1.70+0.031μm) than either the Jacobina (1.50+0.036μm) or Sobral (1.48+0.027μm) colony manifolds which were not significantly different from each other (Fig. 4A).

There was also a highly significant difference in the lengths of the manifolds (Fig. 2) of the 3 types of *L. longipalpis* (df=147; F=116.01; *P*< 0.001). The Campo Grande manifolds (0.94+0.024μm) were significantly longer than the Jacobina manifolds (0.84+0.028μm) which were significantly longer than the Sobral manifolds (0.49+0.012μm) (Fig. 4B).

**Fig. 4.**
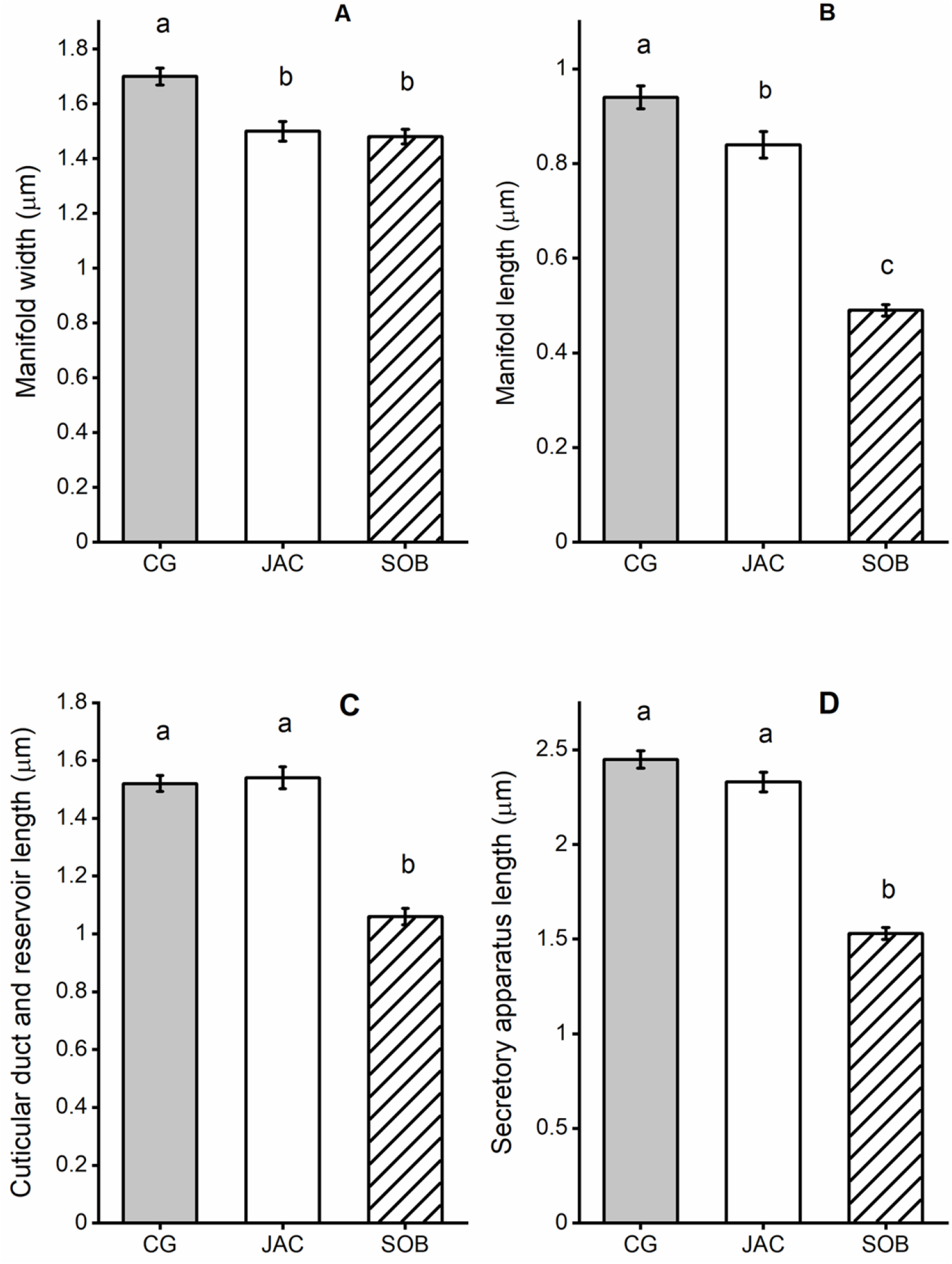
Dimensions of the components of the secretory apparatus observed in 3 members of the *Lutzomyia longipalpis* species complex. Mean size of the measured structures (μm); manifold width (**A)**, manifold length (**B)**, reservoir and cuticular duct length (**C)** and secretory apparatus length (**D)** for each of the three members of the *Lutzomyia longipalpis* species complex; Campo Grande (CG), Jacobina (JAC) and Sobral (SOB). Error bars are ±standard error of the mean. Tukey’s test was used to compare sizes of structures between each member of the complex, measurements with the same letter (a, b or c) were not significantly different (P>0.05) from each other.

There was also a significant difference between the length of the cuticular duct + reservoir (Fig. 2) in the 3 types of *L. longipalpis* (df=147; F=75.55; *P*=0.001). The Campo Grande and Jacobina cuticular ducts + reservoir were not significantly different from each other (1.52+0.027μm and 1.53+0.038μm respectively) whereas the Sobral ducts + reservoir were significantly shorter (1.06+0.028μm) (Fig. 4C).

The overall length of the secretory apparatus (Fig. 2) was also significantly different between the 3 types of *L. longipalpis* (df=147; F=133.53; *P*<0.001). The Campo Grande secretory apparatus was similar in length to the Jacobina secretory apparatus (2.45+0.046μm and 2.33+0.051μm respectively). However, the Sobral secretory apparatus was significantly shorter than either Campo Grande or Jacobina (1.53+0.032μm) (Fig. 4D).

The differences in the size and shape of the secretory apparatus are summarised in Fig. 5. The manifold of the Campo Grande (Fig. 5A) member of the complex was longer and wider than the Jacobina type (Fig. 5C). Overall, the secretory apparatus of the Sobral (Fig. 5B) type was smaller than the others.

**Fig. 5.**
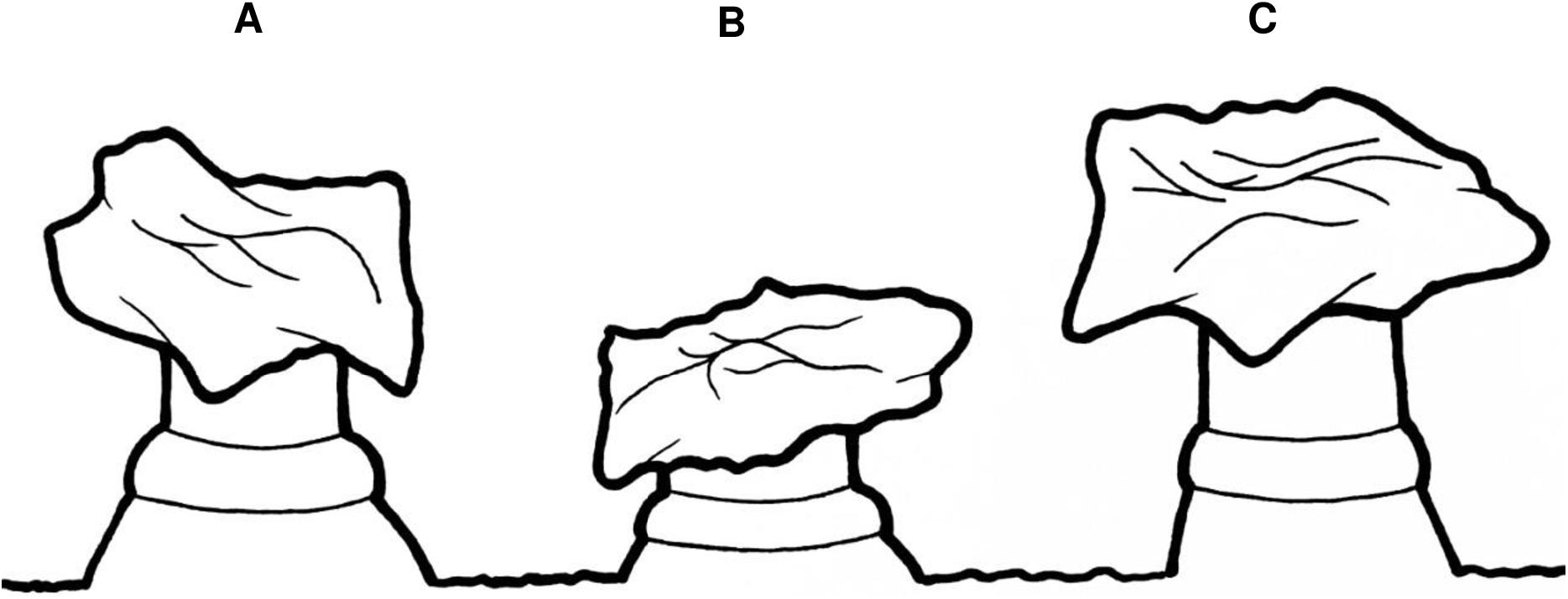
Drawing illustrating the morphological differences observed in the size and shape of the manifold in the three members of the *L. longipalpis* s.l. species complex Campo Grande **(A)**, Sobral **(B)** and Jacobina **(C)**.

### *Migonemyia migonei* secretory apparatus

We found structures resembling the manifold, secretory duct and reservoir (ca. 9/1000mm^2^) previously seen in the *L. longipalpis* in the *M. migonei* samples. These structures were present on the internal cuticle surface of tergites III – VII. This distribution partially matched the distribution of the craters with central pore and spike previously reported on the external surface of tergites III-VI (Costa 2016) (Fig. 6A). The manifold was inserted within a deep recess (av max width 1.50±0.04μm) and appeared to be embedded (ca. 0.25μm deep) within the cuticle. Only the reservoir appeared to be positioned fully within the interior of the abdomen (Figs. 6B and 6C). Multiple observations of the manifolds from different positions suggest that it has the appearance illustrated in Fig. 6 D1 and D2.

**Fig. 6.**
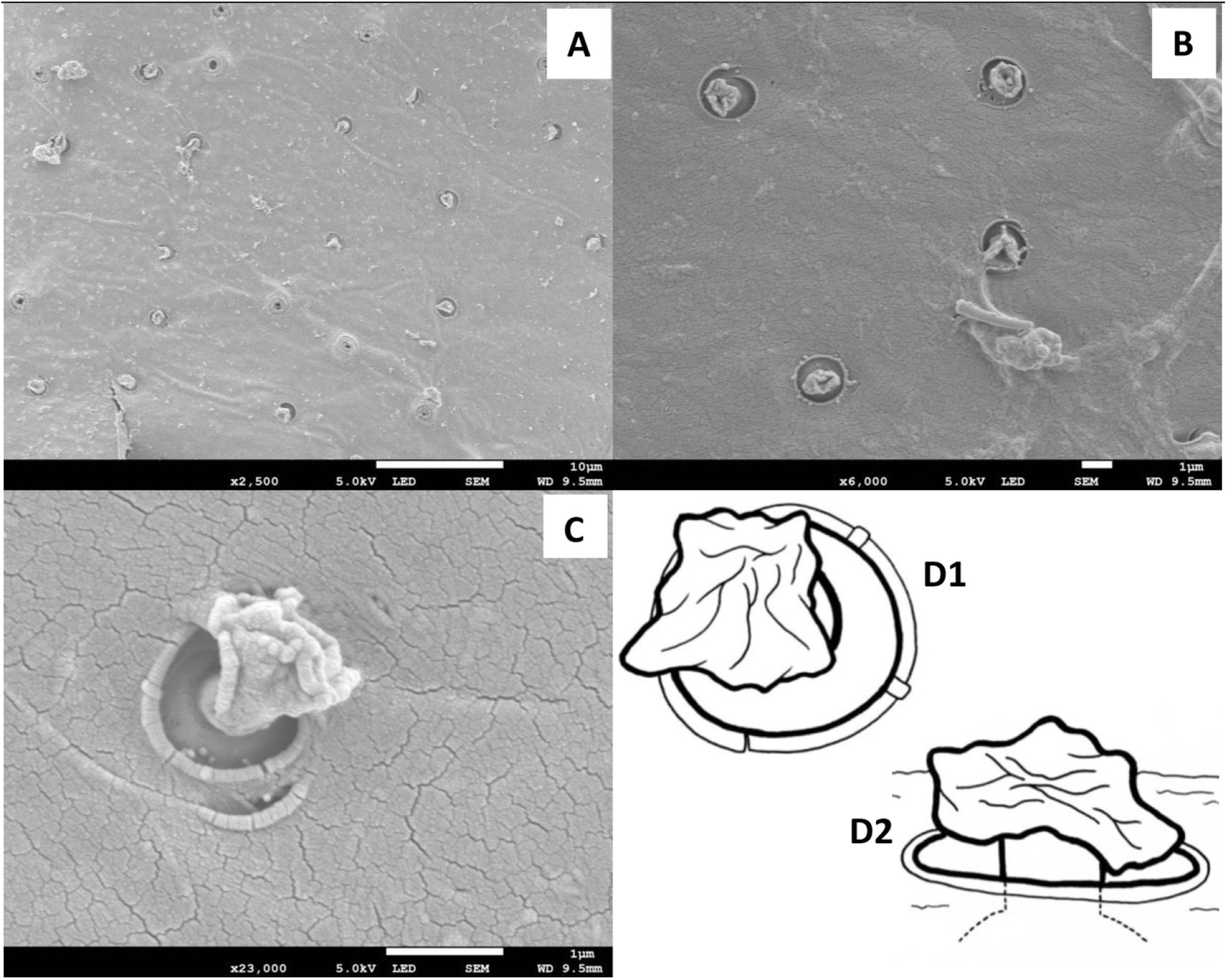
SEM of the interior surface *M. migonei* showing the observable cuticular elements of the secretory apparatus. **A)** Distribution of the secretory structures on the inner surface of the exoskeleton on tergite III, **B)** secretory apparatus set within a deep pocket embedded in the exoskeleton on tergite IV, **C)** close-up of a secretory unit showing the manifold embedded within the cuticle observed at the bottom of the pocket on tergite III. **D1)** Drawing of the *M. migonei* secretory apparatus from above (top left) showing the reservoir positioned over the hole in the cuticle and then **D2)** a side-on view showing the reservoir connected via secretory tubule to the top of the manifold sitting within the hole in the cuticle.

## Discussion

Pheromone disseminating structures have been observed on the cuticle of 53 species of New World *(Lutzomyia* and *Brumptomyia* spp) and 5 Old World *(Sergentomyia)* species (Ward *et al.* 1991; Ward *et al.* 1993). These structures take a diverse range of morphological forms and include structures such as pores in craters, pores with emergent spines, mammiform papules with or without spines and apple shaped structures (Ward *et al.* 1993). This study reveals that in addition to the pheromone disseminating structures visible on the external surface of the abdomen there are additional cuticular structures on the inside surface of the abdominal cuticle which have not been observed or described before. Each external structure is associated with a new structure which we have called the manifold (a device used to aggregate or distribute gases or fluids). Although the precise function of the manifold is unknown it is connected via a tubule to the end apparatus of the secretory apparatus thus it is clearly associated with the distribution of the pheromone from the secretory apparatus to the external surface of the sand fly.

SEM has been widely used to examine the external secretory apparatus and other externally visible cuticular structures in sand flies and other insects. The cells associated with pheromone production have been examined by TEM in sand flies (Costa 2016; Lane and S. 1990; Spiegel *et al.* 2011; Spiegel *et al.* 2002; Ward *et al.* 1993) and other pheromone producing insect groups e.g. Lepidoptera, Coleoptera, Hymenoptera and species of Trichoptera from the families Rhyacophilidae and Limnephilidae (Lensky *et al.* 1985; Melnitsky and Deev 2009; Nardi *et al.* 1996; Noirot and Quennedey 1974; Noirot and Quennedey 1991; Percy 1979; Percy 1975; Pierre *et al.* 1996; Raina *et al.* 2000). Most of these studies were carried out to describe the arrangement, location and/or distribution of the pheromone gland secretory cells they were not carried out to examine the mechanisms by which the pheromone was transported from the site of biosynthesis to the point of dissemination on the surface of the cuticle. We are not aware of any published SEM studies that have examined the internal structures associated with pheromone production and transport in sand flies or any other group of insects.

In this study we used *L. longipalpis* from Lancaster University that had been stored in hexane and *M. migonei* from Charles University colony that had been stored in 70% ethanol. Samples stored in ethanol required longer KOH digestion to remove the interior abdominal tissue. We found digestions up to four hours useful for *L. longipalpis* specimens and up to 10 hours for *M. migonei* specimens.

In addition to the differences in the structure of their sex-aggregation pheromone, the members of the *L. longipalpis* s.l. species complex analysed in this study, have also shown differences related to the biosynthesis and release of their pheromones (Gonzalez *et al.* 2017). The results of this study also show significant morphological differences between the size and shape of the manifolds. Interestingly there have been no reported differences in the size and shape of the papules which can be observed on the surface of the tergites in *L. longipalpis* s.l. The manifolds and other elements of the secretory apparatus of the Sobral member of the complex are significantly shorter than either Campo Grande or Jacobina. The manifold width of Sobral *L. longipalpis* is not significantly different to that of Jacobina but both are significantly narrower than in Campo Grande. The overall effect of the differences is that the Jacobina and Campo Grande are similar in size and shape to each other whereas the Sobral structure appears shorter and squatter. The effect of these differences may be to position the secretory cells that would surround the end apparatus closer to the surface in the Sobral type than the other 2 types. This may reflect the difference in the molecular weight of the 2 methylsesquiterpenes (m.w. 218) found in Campo Grande and Jacobina compared to the molecular weight of the diterpene pheromone (m.w. 272) in the Sobral population. Thus, the distance for the larger molecule to travel from the secretory cell to the external surface is less than for the other two lighter and less volatile molecules.

The manifold of *M. migonei* is very different to those observed in *L. longipalpis* s.l. and is positioned within the tergal cuticle in a pit-like structure. The end apparatus is connected by a short duct to the manifold. The effect of this arrangement is that the secretory cells would be much closer to the surface than in *L. longipalpis* and this may reflect a relatively lower volatility (either higher molecular weight or presence of functional groups) of any sex aggregation pheromone produced by *M. migonei.* Although there is behavioural evidence for the presence of a sex-aggregation pheromone in *M. migonei* no compound(s) with a similar chemical profile to the sex aggregation pheromones found in the *L. longipalpis* s.l. species complex has been found (Costa 2016).

The density of manifolds found on the internal cuticle of the sobralene producing Sobral (CE) *L. longipalpis* was 18 per 1000 μm^2^ (ca. 3469 in total) and matched the density of papules previously observed on the tergal surface of *L. longipalpis* from Sobral, (19 per 1000 μm^2^) (Spiegel *et al.* 2002). This is not dissimilar to estimates of 14 per 1000 μm^2^ for the same Sobral population (Lane and Ward 1984). The density of manifolds in the Campo Grande (MS) (*S*)-9-methylgermacrene-B producing population was approximately 13 per 1000 μm^2^, part-way between the 8 per 1000 μm^2^ papules observed by Lane and Ward (1984) in *L. longipalpis* collected at Lapinha Cave (MG) and 21 per 1000 μm^2^ papules in *L. longipalpis* also collected at Lapinha Cave (Spiegel *et al.* 2002). The meaning of this difference is unclear, it may be related to significant differences between the Campo Grande population and the Lapinha population similar to those observed between the Sobral (*S*)-9-methylgermacrene-B and the Lapinha population in which the Sobral population was found to produce significantly more pheromone than the Lapinha population (Hamilton *et al.* 2005) and principal component analysis of SNPs in 245 chemoreceptor genes (Hickner *et al.* 2021).

This is the first time that the manifold structure has been seen in any group of insects and its function is unclear. It may be that the manifold is only found in Phlebotomine sand flies, but it may occur in other insect orders. It could simply be a device to ensure the safe transport of the sex aggregation pheromone from the secretory cells through the cuticle. The sturdiness of the structure could suggest that it is designed to minimise potential leakage of the potentially toxic terpene (Agus 2021) pheromone into the abdomen. Male sand flies engage in combat with other males to defend territory and in these aggressive battles (Jarvis and Rutledge 1992; Soares and Turco 2003) males could potentially risk dislodging unprotected plumbing carrying toxic pheromone. However, without a clear view of the interior of the manifold it is uncertain if additional functionality may exist e.g. a passive or controllable valve or a reservoir of pheromone or other mechanism to regulate pheromone flow to help provide a supply of pheromone when it is required (Gonzalez *et al.* 2017). In the future it may be possible to get a clear view of the interior of these structures using Synchrotron Radiation Microtomography (Enriquez *et al.* 2021).

It was difficult to count the papules on the external abdominal cuticle because of the presence of macrotrichia and other structures (Lane and Ward 1984; Spiegel *et al.* 2002). Observing the location, distribution and density of the manifolds on the inner cuticle was a convenient way to check the whole inner cuticle of the abdomen for secretory devices. More studies should now be conducted to compare the number of these structures in different members of the *L. longipalpis* s.l. complex and from different parts of Brazil as well as to determine their distribution in other New and Old-World species.

The *M. migonei* manifold lay within the cuticle and although it was possible to observe it within a clearly defined hole we could not check morphological details. The details of how the secretory apparatus is connected to the exterior remains elusive and although there was one manifold per hole it was not possible to clarify if there was more than one opening per pheromone secreting structure (“spined crater” (Ward *et al.* 1993)) on the exterior of the insect. We found that the manifolds were distributed on tergites III to VII but more studies should be carried to fully describe the morphology of the manifold and then link the morphological form to the pheromone and its function.

These results may contribute to the discussion of the nature of the *L. longipalpis* species complex, as they show that there are clear morphological differences between 3 of the members of the complex. These structures may also be useful taxonomic tools more generally within the Phlebotominae. The study also shows that in addition to the widespread distribution of the external structures linked to pheromone production these internal structures are likely to be strongly associated with active pheromone production and therefore their presence in species where pheromone production has been inferred through behavioural studies but not confirmed through chemical analysis should be undertaken. The presence of the manifold and its associated end apparatus is considerably easier to locate than hidden isolated external structures as in *L. renei* (Spiegel *et al.* 2002) therefore locating the secretory apparatus and thus identifying which sand fly species may produce pheromone will be easier. Behavioural analysis in the lab and field has shown that female *Phlebomomus papatasi* and *P. argentipes* are attracted to conspecific males, however no external structure has been observed on the abdomen. This approach may simplify the search for the pheromone source.

## Acknowledgements

We are grateful to Lisa Butler for assistance with the sand fly colonies at Lancaster University and Tomas Becvar for breeding the colony of *L. migonei* at Charles University.

To Dr. Sara Baldock in Lancaster University Chemistry Department for the help with acquisition of SEM images and to Dr Nigel Fullwood, Lancaster University, for useful discussions on EM.

## Funding Statement

This study was financed in part by the Coordenação de Aperfeiçoamento de Pessoal de Nível Superior – Brasil (CAPES) – Finance Code 001 and Fundação de Amparo à Pesquisa do Estado de Minas Gerais (FAPEMIG - PPM-00792-18). GBT was supported by CAPES/PRINT (41/2017). JDAF received a research fellowship from “Conselho Nacional de Desenvolvimento Científico e Tecnológico” (CNPq - 303680/2020-2). PV was supported by ERDF funds CZ.02.1.01/0.0/0.0/16 019/0000759. JGCH was supported by The Wellcome Trust (080961/Z/06/Z).

The funders had no role in study design, data collection and analysis, decision to publish, or preparation of the manuscript.

## Consent for publication

Not applicable

## Availability of data and materials

All data generated or analysed during this study are included in this published article.

## Competing interests

The authors have declared that no competing interests exist.

## CRediT author statement

**Gabriel B. Tonelli**: Methodology, Investigation, Original Draft, Writing-Review & Editing, Visualization.

**J.D.A. Filho**: Resources, Writing-Original Draft, Project administration, Funding acquisition

**A. M. Campos:** Formal analysis, Writing – Review & Editing

**C.M. de Souza:** Investigation, Writing – Review & Editing

**A.R. Amaral:** Writing – Review & Editing, Visualization

**P. Volf:** Methodology, Resources, Writing - Review & Editing

**E. Shaw:** Methodology, Resources, Writing – Review & Editing

**Gordon Hamilton:** Conceptualization, Methodology, Resources, Writing-Original

Draft, Writing-Review & Editing, Visualization, Project administration, Funding acquisition.

## Notes

### Competing Interest Statement

The authors have declared no competing interest.

